# Fc-modified SARS-CoV-2 neutralizing antibodies with therapeutic effects in two animal models

**DOI:** 10.1101/2022.06.21.496751

**Authors:** Masaru Takeshita, Hidehiro Fukuyama, Katsuhiko Kamada, Takehisa Matsumoto, Chieko Makino-Okamura, Tomomi Uchikubo-Kamo, Yuri Tomabechi, Kazuharu Hanada, Saya Moriyama, Yoshimasa Takahashi, Hirohito Ishigaki, Misako Nakayama, Cong Thanh Nguyen, Yoshinori Kitagawa, Yasushi Itoh, Masaki Imai, Tadashi Maemura, Yuri Furusawa, Hiroshi Ueki, Kiyoko Iwatsuki-Horimoto, Mutsumi Ito, Seiya Yamayoshi, Yoshihiro Kawaoka, Mikako Shirouzu, Makoto Ishii, Hideyuki Saya, Yasushi Kondo, Yuko Kaneko, Katsuya Suzuki, Koichi Fukunaga, Tsutomu Takeuchi, the Keio Donner Project

## Abstract

The use of therapeutic neutralizing antibodies against SARS-CoV-2 infection has been highly effective. However, there remain few practical antibodies against viruses that are acquiring mutations. In this study, we created 494 monoclonal antibodies from COVID-19–convalescent patients, and identified antibodies that exhibited comparable neutralizing ability to clinically used antibodies in the neutralization assay using pseudovirus and authentic virus including variants of concerns. These antibodies have different profiles against various mutations, which were confirmed by cell-based assay and cryo-electron microscopy. To prevent antibody-dependent enhancement, N297A modification was introduced, and showed a reduction of lung viral RNAs by therapeutic administration in a hamster model. In addition, an antibody cocktail consisting of three antibodies was also administered therapeutically to a macaque model, which resulted in reduced viral titers of swabs and lungs and reduced lung tissue damage scores. These results showed that our antibodies have sufficient antiviral activity as therapeutic candidates.

## Introduction

Severe acute respiratory syndrome coronavirus 2 (SARS-CoV-2) continues to spread with the acquisition of various mutations. In Japan, the original Wuhan strain acquired the D614G mutation in the early stages (Korber et al., 2020), and was replaced by the more infectious Alpha variant from the end of 2020 to the first half of 2021 (Volz et al., 2021). Next, it was replaced by the much more infectious Delta variant (Earnest et al., 2021), and then further replaced by the Omicron variant as of November 2021 (Shah and Woo, 2022). Various vaccines have been developed against the Wuhan strains and, fortunately, they have shown efficacy against variants (Thomas et al., 2021). Although the number of cases has decreased in some countries, probably due to vaccine effectiveness, the global pandemic has not yet been terminated.

The novel coronavirus disease 2019 (COVID-19) is known to progress to a severe state due to excessive immune response and inflammation in the late stages of the disease (Siddiqi and Mehra, 2020). Therefore, immunosuppressants, such as steroids, IL-6 inhibitors, and JAK inhibitors, are used in this stage (Lowery et al., 2021; Yokota et al., 2021). In contrast, in the early stage of infection, there is a period of viral replication, and antiviral therapy is effective during this period. The development of antibody therapies is progressing rapidly, with the Food and Drug Administration (FDA) granting emergency use authorization (EUA) for Regeneron’s antibody cocktail (casirivimab and imdevimab), Lilly’s antibody (bamlanivimab), the cocktail of bamlanivimab and etesevimab, GSK’s sotrovimab, the cocktail of tixagevimab and cilgavimab for prevention, and most recently bebtelovimab monotherapy. These therapies decrease the risk of hospitalization and death by 70% to 85% (Dougan et al., 2021; Weinreich et al., 2021; Gupta et al., 2021), however, the EUA for tixagevimab and cilgavimab and bebtelovimab monotherapy has been revoked because of decreased efficacy against newly emerged variant of concerns (VOCs). There remain a limited number of treatment options.

We have been collecting peripheral blood samples from convalescent patients since the beginning of the COVID-19 epidemic in Japan in March 2020, from which we started to develop neutralizing antibodies. We have identified several antibodies with neutralizing ability that is equivalent to therapeutic antibodies, and we have demonstrated their efficacy by pseudovirus and authentic virus neutralization assay *in vitro*, and by infection experiments with hamster and macaque models *in vivo*.

## Results

### Selection of patients with high neutralizing antibody titer

We collected peripheral blood samples from COVID-19 patients who were discharged from Keio University Hospital. Patient characteristics are shown in Supplementary Table 1. The neutralization ability of sera was analyzed by cell-based Spike-ACE2 inhibition assay (Figure 1A). From them, we selected 12 patients with high neutralizing titer for antibody production (Figure 1B). Their characteristics are shown in Supplementary Table 2. From the peripheral blood B cells of these patients, we sorted RBD and S1-binding memory B cells and antigen-nonspecific plasma cells (Figure 1C). The sequences of H-chain and L-chain variable regions were amplified by polymerase chain reaction (PCR), and inserted into expression vectors to produce monoclonal antibodies. A total of 494 antibodies were produced, 408 from antigen-specific memory B cells, and 86 from antigen-nonspecific plasma cells.

**Figure 1.**
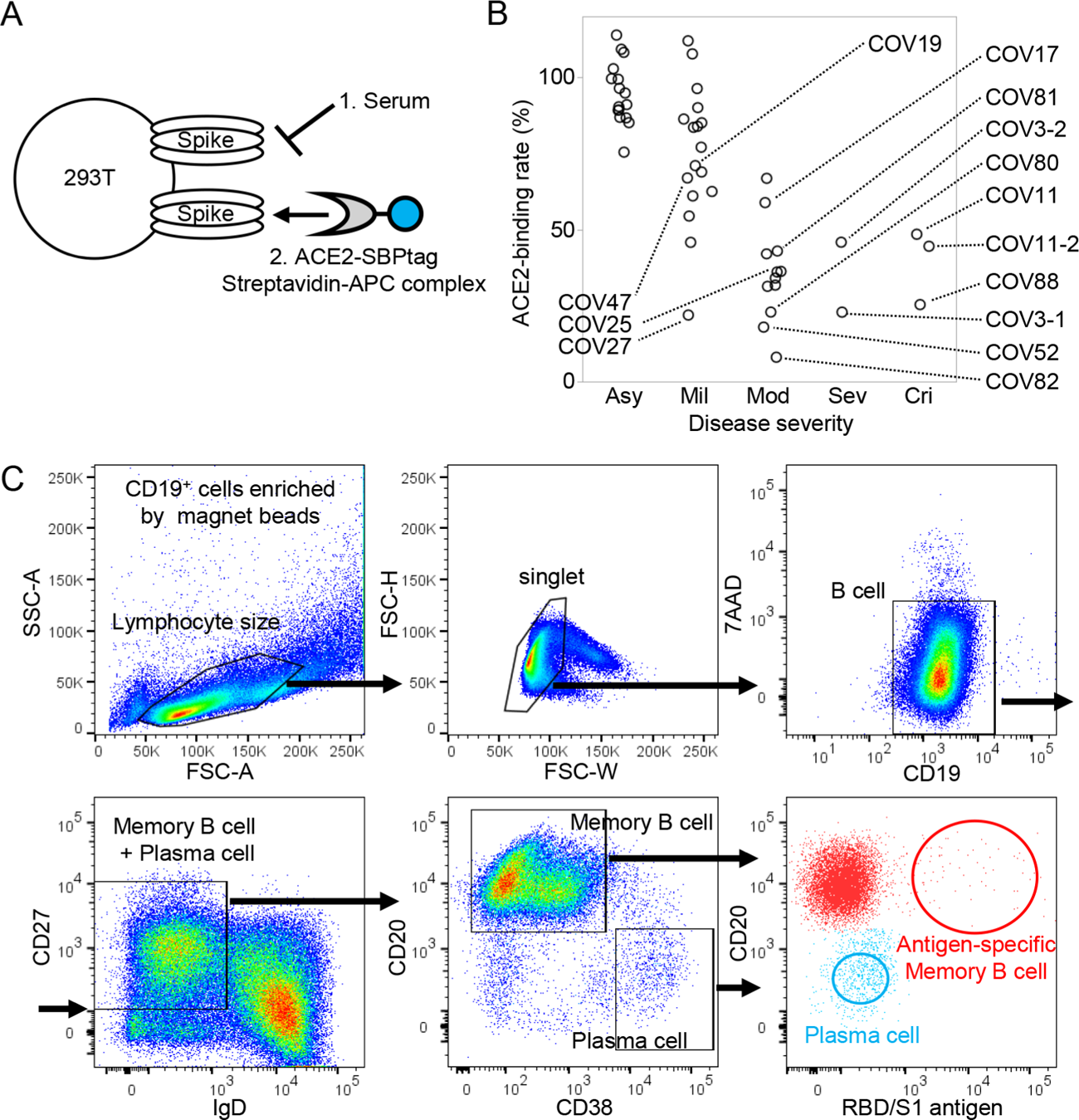
Patient selection and cell sorting Serum neutralization titers of 47 COVID-19 convalescent patients were measured by (A) cell-based Spike-ACE2 inhibition assay. The binding quantity of soluble ACE2 to Spike-expressing cells without serum/antibody is defined as 100%, and the binding quantities of soluble ACE2 to Spike-expressing cells after incubation with serum/antibody are calculated as the ACE2-binding rate. (B) The neutralization ability of patient serum for each severity is shown. Samples used for antibody production are labeled with their ID. (C) The sorting strategy is shown. CD19^+^ cells were size gated, and CD19^+^CD27^+^IgD^-^ cells were selected. Antigen-specific memory B cells (red) and antigen-nonspecific plasma cells (blue) were single-cell sorted. Asy, asymptomatic; Mil, mild; Mod, moderate; Sev, severe; Cri, critical.

### Screening of neutralizing antibodies

We screened patient-derived antibodies using two procedures. The first was the Spike-ACE2 inhibition assay described above, in which monoclonal antibodies were used instead of serum. As shown in Figure 2A, although the majority of the antibodies clustered around 100% of the ACE2-binding rate, indicating that they did not compete with ACE2, part of the antibodies from memory B cells and two antibodies from plasma cells showed inhibition of ACE2 binding. The peripheral blood of COV003 and COV011 was taken twice, at day 35 and 109, and at day 42 and 63, after the first positive PCR, respectively, and neutralizing antibodies were mainly found in the later samples.

**Figure 2.**
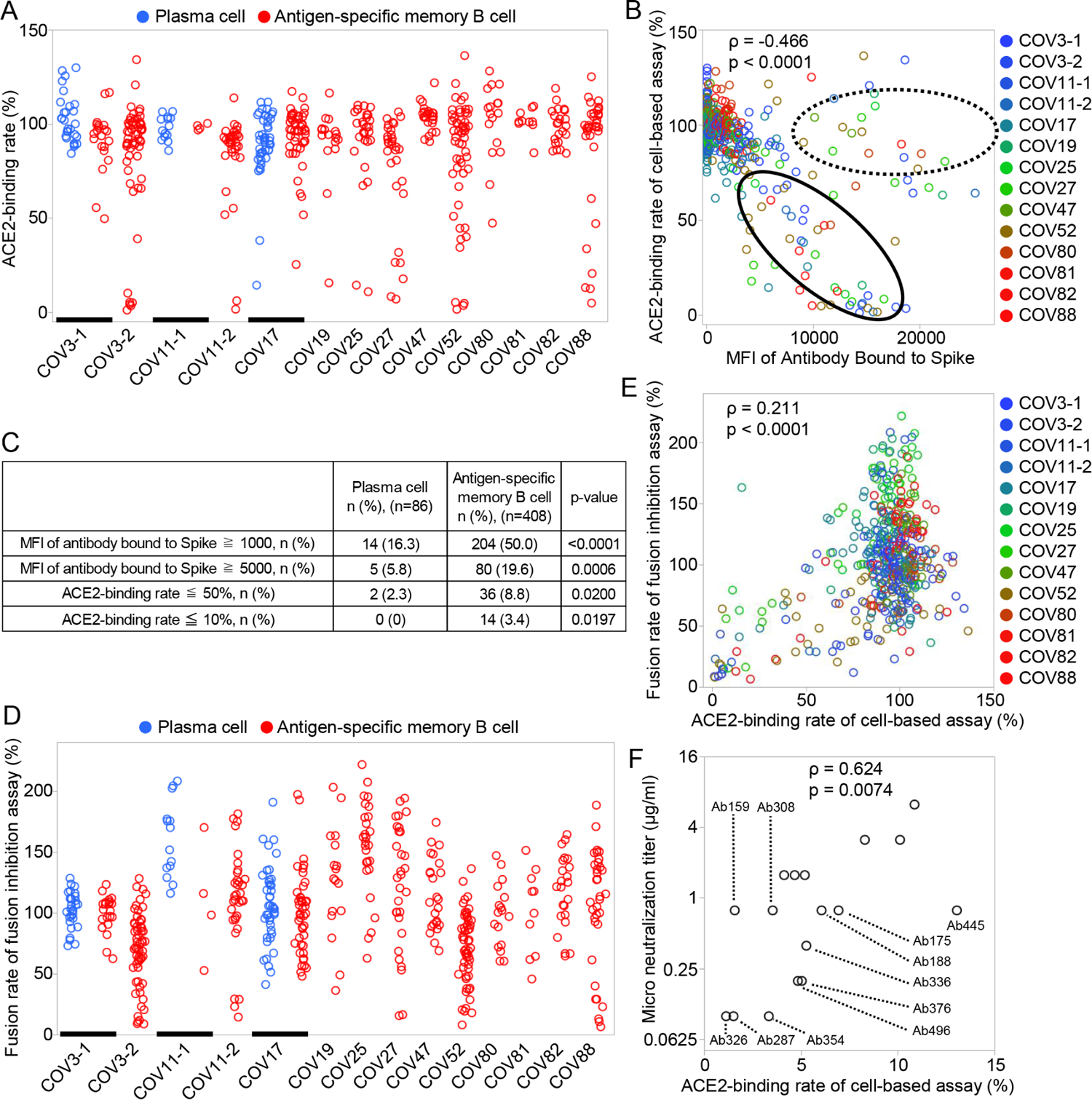
Screening of neutralizing antibodies (A) Neutralization ability of recombinant monoclonal antibody was measured by the cell-based Spike-ACE2 inhibition assay. Blue indicates plasma cell–derived antibody, and red indicates antigen-specific memory B cell–derived antibody. (B) Negative correlation of the ACE2-binding rate and the median fluorescent intensity (MFI) of antibodies bound to Spike-expressing cells is shown. Antibodies that have binding ability without neutralization ability are encircled by a dotted line, and antibodies that have correlation between binding and neutralization ability are encircled by a solid line. Spearman’s rank correlation coefficient was calculated from all samples. (C) The percentage of antibodies having high binding ability to Spike and high ability to inhibit ACE2 binding to Spike is shown by the source cell. (D) Neutralization ability of recombinant monoclonal antibody was measured by fusion inhibition assay. The quantity of cell fusion without antibody is defined as 100%, and the quantity of cell fusion after incubation with antibodies is shown as the fusion rate. Blue indicates plasma cell–derived antibody, and red indicates antigen-specific memory B cell–derived antibody. (E) The correlation between the ACE2-binding rate and the fusion rate of each antibody is shown. Spearman’s rank correlation coefficient. (F) The correlations between the end-point micro neutralization titer and the ACE2-binding rate are shown. Spearman’s rank correlation coefficient.

Next, we examined the relationship between the binding ability of these antibodies against Spike-expressing cells and their ability to inhibit the binding of ACE2 against Spike-expressing cells. As shown in Figure 2B, these antibodies were divided into two groups: antibodies that have binding ability without neutralization, as shown by the dotted line, and antibodies that have binding ability correlated with neutralization ability, as shown by the solid line. Figure 2C shows the proportions of antibodies that bind to and neutralize Spike by source of B cell type. Although over 80% of the antibodies produced from antigen-nonspecific plasma cells did not neutralize or even bind to Spike-expressing cells, approximately half of the antigen-specific memory B cell–derived antibodies could bind to Spike, 20% bound strongly, 9% had neutralizing ability, and 3.4% had high neutralizing ability.

In order to perform screening of these antibodies more robustly, we also examined the neutralizing ability by cell fusion assay, which examines the extent to which antibodies inhibit the fusion of Spike-expressing cells and ACE2-expressing cells. As shown in Figure 2D, the antibodies with high neutralizing ability were detected to be similar to those in the cell-based Spike-ACE2 inhibition assay. The results of these two procedures were also correlated (Figure 2E).

In order to confirm that the selected antibodies have the ability to neutralize the authentic virus, we performed end-point micro neutralization assay to determine the minimum concentration required to neutralize authentic viruses for top antibodies. As shown in Figure 2F, the micro neutralization titers and ACE2-binding rates were well-correlated, and 11 antibodies were found to be able to completely neutralize authentic virus at a concentration of less than 1 µg/ml.

### Affinity and epitope binning by Biolayer interferometry

To further characterize candidate antibodies, we measured the affinity against RBD protein and examined the epitope overlapping. As shown in Supplementary Figure 1A, selected antibodies have a low KD value of 1 × 10^-9^ M to 1 ×10^-12^ M. The KD value did not correlate with the minimum neutralization titer (Supplementary Figure 1B). We also performed epitope binning assay of the top candidates, however, all of the top candidate antibodies had overlapping of their epitopes (Supplementary Figure 1C).

### The effect of point mutation of Spike on the neutralizing ability of antibodies

Next, we investigated how the selected antibodies were affected by various mutations in the cell-based Spike-ACE2 inhibition assay using mutated Spike-expressing cells. The ACE2-binding rates of each antibody for mutations within and outside the RBD are shown in Figure 3A and 3B, respectively. Each antibody showed variable neutralizing ability at various sites within the RBD, and these amino acids were considered to be candidates of epitopes. Among them, the E484K mutation affected at least 8 of the top 11 antibodies, and mutation at W406, K417, F456, T478, F486, F490, and Q493 affected 3 to 4 of 11 antibodies. These facts indicate that these positions may be major epitopes of human humoral immunity against Wuhan-hu-1 strain, consistent with previous reports (Hoffmann et al., 2021; Wang et al., 2021b; Zhou et al., 2021). Because these antibodies were derived from B cells that bind to RBD/S1 and are selected by ACE2 inhibition, it is possible that they were not significantly affected by mutations outside the RBD, including the N-terminal domain (Figure 3B). We also examined the cells expressing Spike, including all variant mutations of SARS-CoV-2 and SARS-CoV-1 (Figure 3C). Although efficacy varies from strain to strain, the Omicron (BA.1) variant has become resistant to almost all antibodies except for Ab188. In addition, the neutralizing ability of any antibodies examined was not observed against SARS-CoV-1.

**Figure 3.**
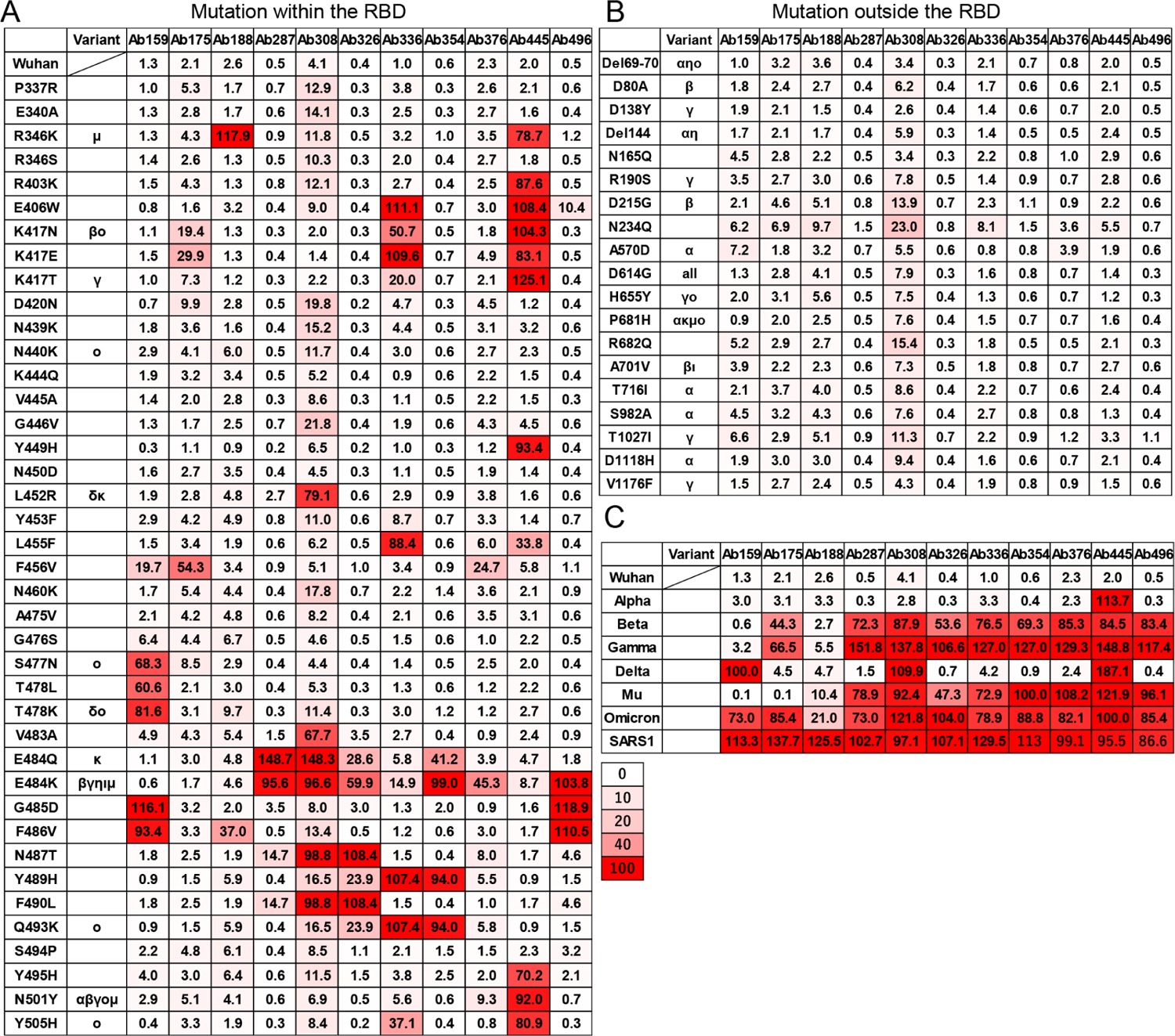
Effect of point mutation of Spike protein on antibody neutralizing ability The ACE2-binding rate (%) of recombinant monoclonal antibodies against Spike protein with various point mutations (A) within RBD, (B) outside RBD, and (C) against Spike proteins of VOCs and SARS-CoV-1 were measured by the cell-based Spike-ACE2 inhibition assay. The variant column indicates that the mutation is contained in the VOC and Variants of interest. The numbers indicate the binding quantities of soluble ACE2 to Spike-expressing cells after incubation with antibodies. The color indicates the grade of neutralization ability.

### Pseudovirus and authentic virus neutralization assay

Before variants of concern (VOCs) appeared, we selected five antibodies, and performed neutralization assay using pseudovirus with Spike protein of the original Wuhan strain and four major variants. The IC_50_ of each antibody was measured comparing with the therapeutic antibody imdevimab as a control (Figure 4AB). Ab159, Ab326, Ab354, and Ab496 exhibited similar or better neutralizing ability against Wuhan pseudovirus than the therapeutic antibody. However, Ab159 could not neutralize the Delta variant due to T478K mutation, whereas Ab326, Ab354, and Ab496 could not neutralize the Beta and Gamma variants due to E484K mutation (Figure 3A).

**Figure 4.**
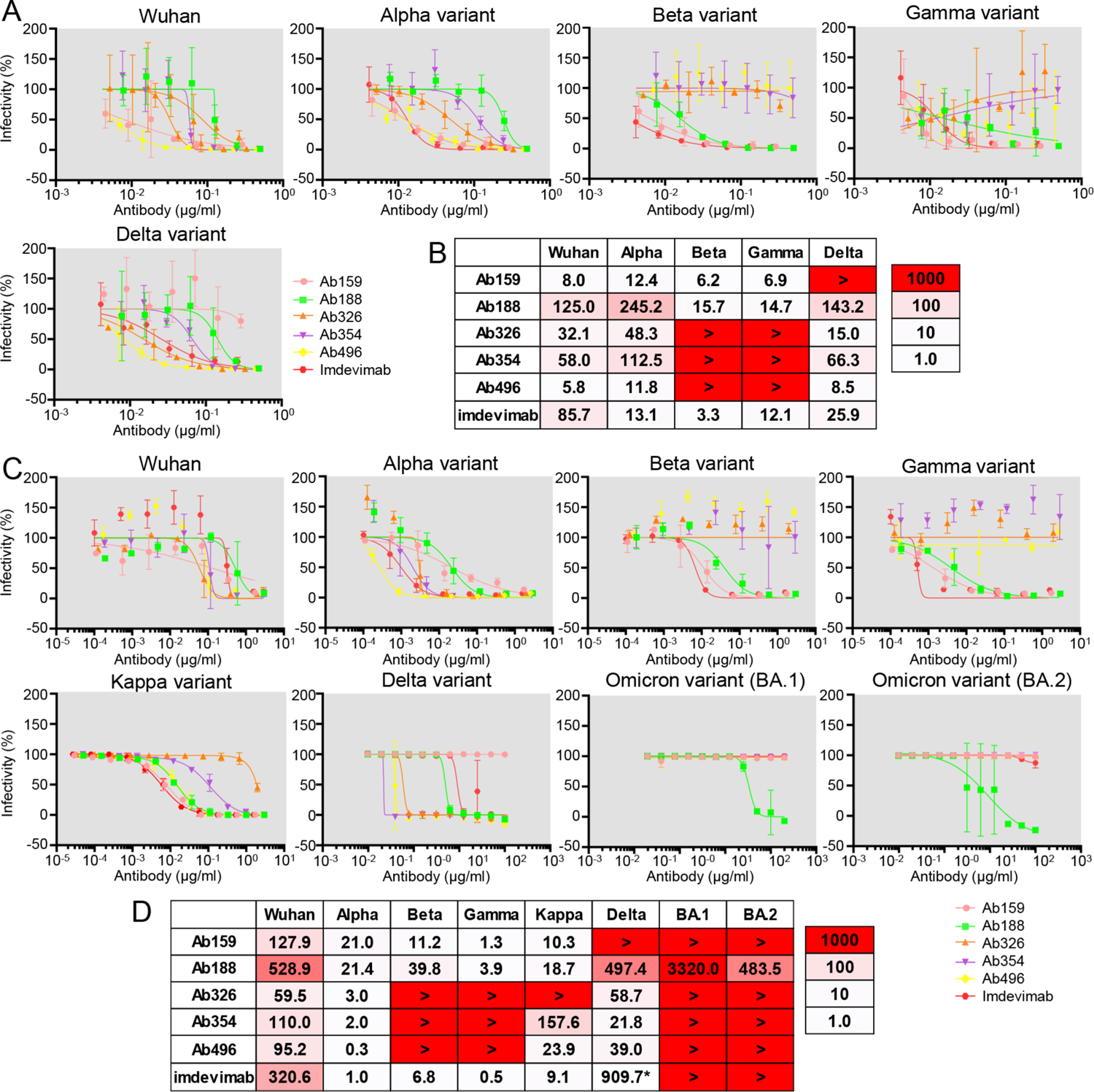
Pseudovirus and authentic virus neutralization assay. The IC_50_ (ng/ml) of selected antibodies was measured using pseudovirus that expresses Spike of the Wuhan strain or Alpha, Beta, Gamma, or Delta variants. The data are represented as mean ± SD. (A) The neutralization curve and (B) IC_50_ values are shown. The color indicates the neutralizing ability. The IC_50_ (ng/ml) of the selected antibodies was examined using authentic virus of the Wuhan strain or Alpha, Beta, Gamma, Delta, Kappa, Omicron (BA.1 and BA.2) variants. The data are represented as mean ± SD. (C) Neutralization curve and (D) IC_50_ values are shown. *Measured separately.

After VOCs appeared, we examined the neutralization abilities of antibodies against authentic virus of WK-521 and variants, including Alpha, Beta, Gamma, Delta, Kappa, and Omicron BA.1 and BA.2 (Figure 4CD). Similar to the results of pseudovirus, Ab326 and Ab354 were ineffective against the Beta and Gamma variants, and Ab159 was ineffective against the Delta variant. Whereas many antibodies lost neutralizing ability against both Omicron variants, Ab188 retained neutralizing ability. Although the IC_50_ of imdevimab against the Wuhan strain was reported to be 6 to 70 ng/ml (Fenwick et al., 2021; Hansen et al., 2020; Wang et al., 2021a), our experiments resulted in 320.6 ng/ml, which is higher than previously reported, probably due to the differences in experimental conditions, such as the usage of a sensitive cell-line, VeroE6 cells expressing TMPRESS2.

### Cryo-electron microscopy

To gain structural insights into antibodies and the SARS-CoV-2 spike, we performed cryo-electron microscopy (cryo-EM) analysis of complexes with the proline-substituted stable spike (Supplementary Table 3). In addition to the five selected antibodies, Ab445 was also included in the analysis, as it was assumed to have a different epitope from the others (Figure 3). Single-particle analysis of a series of complexes yielded a better quality map of a class with the most abundant particles. However, all of the reconstructed densities of the interface between RBD and Fab were poor, probably due to the individual motion of RBD within the spike trimer. Accordingly, we performed local refinement to improve the density for each Fab-RBD portion, which allows fine modeling of the variable domain of Fab bound to RBD.

In this analysis, the Fab binding location to RBD is roughly classified into three locations (Figure 5A). First, class I type (Barnes et al., 2020) Fabs (Ab188, Ab445) were observed, which access from the same side as the ACE2-binding surface on RBD. Ab326, Ab354, and Ab496 belong to the Class II type antibody that binds from the upper side of RBD. In particular, an *N*-linked glycan was identified at Asn59 of the heavy chain of Ab354 (Supplementary Figure 2). The fucose moiety of the glycan significantly contributes to binding on the loop region, as previously reported as COVOX-316 (Dejnirattisai et al., 2021). A fairly large number of the buried solvent-accessible area per RBD (∼1850 / ∼1520 Å^2^, +sugar / -sugar) is occupied for this interaction, and each Fab approaches the root of a neighbor RBD. The circular inclination of bound Fab with tight binding might be effective to prevent it from lifting RBD (Supplementary Figure 2).

**Figure 5.**
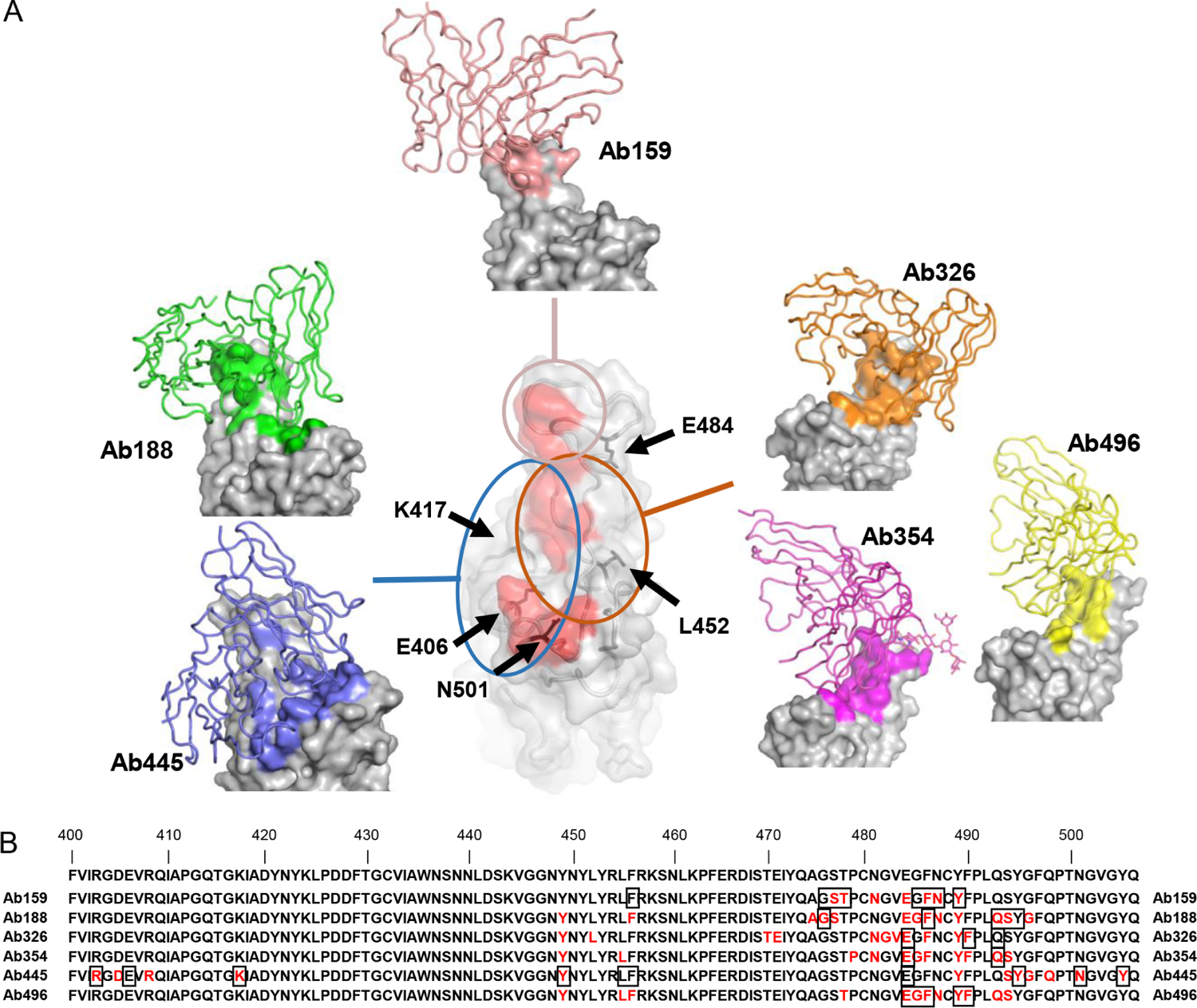
Cryo-EM structure of neutralizing antibodies (A) The structures of RBD and Ab159, Ab188, Ab326, Ab354, Ab445, and Ab496 are shown. Only the variable domains of antibodies are modeled and drawn as a cartoon tube (individual color) on the RBD surface (gray), and the epitope of each antibody is colored the same as each antibody. The red area in the central RBD is the binding residue of ACE2 (7A94) (Benton et al., 2020), showing the relationship between the binding sites of the antibodies, which are roughly divided into three groups. The positions of key amino acids are indicated by black arrows. (B) The residues 400-506 of Spike are shown. The epitopes revealed by cryo-EM are colored in red, and the residues affected by the mutation described in Figure 3A are shown in squares.

The third group, Ab159, exclusively binds with the major loop of RBD. Unlike the recently reported antibodies, A23-58.1, B1-182.1 (Wang et al., 2021a), and COVOX-253 (Dejnirattisai et al., 2021), Ab159 limitedly recognizes an apex of the loop, and mainly interacts only through CDR3 of the heavy chain. The total buried solvent-accessible area is ∼1180 Å^2^, the smallest area among the analyzed structures (Figure 5A).

We defined the epitope as residues that interact with each other within 4Å of distance between their respective central atom. Epitopes of each antibody are shown in red in Figure 5B and were localized around residues 470-500 of Spike. The residues that affected the neutralizing ability in the Spike-ACE2 inhibition assay described in Figure 3A are marked with squares, and are highly consistent with the results of the structural analysis.

### Fc-engineering to prevent antibody-dependent enhancement

In order to prevent antibody-dependent enhancement (ADE), our antibodies used in *in vivo* study had the N297A mutation introduced in the IgG1-Fc region, which reduces binding to the Fc receptor (Chao et al., 2009). We checked Fc-mediated antibody uptake before and after the introduction of the N297A mutation using Raji cells. As shown in Figure 6, the antibody without N297A showed Fc-mediated antibody uptake in the concentration range of 1 to 10 μg/ml whereas the uptake was almost abolished by the introduction of N297A.

**Figure 6.**
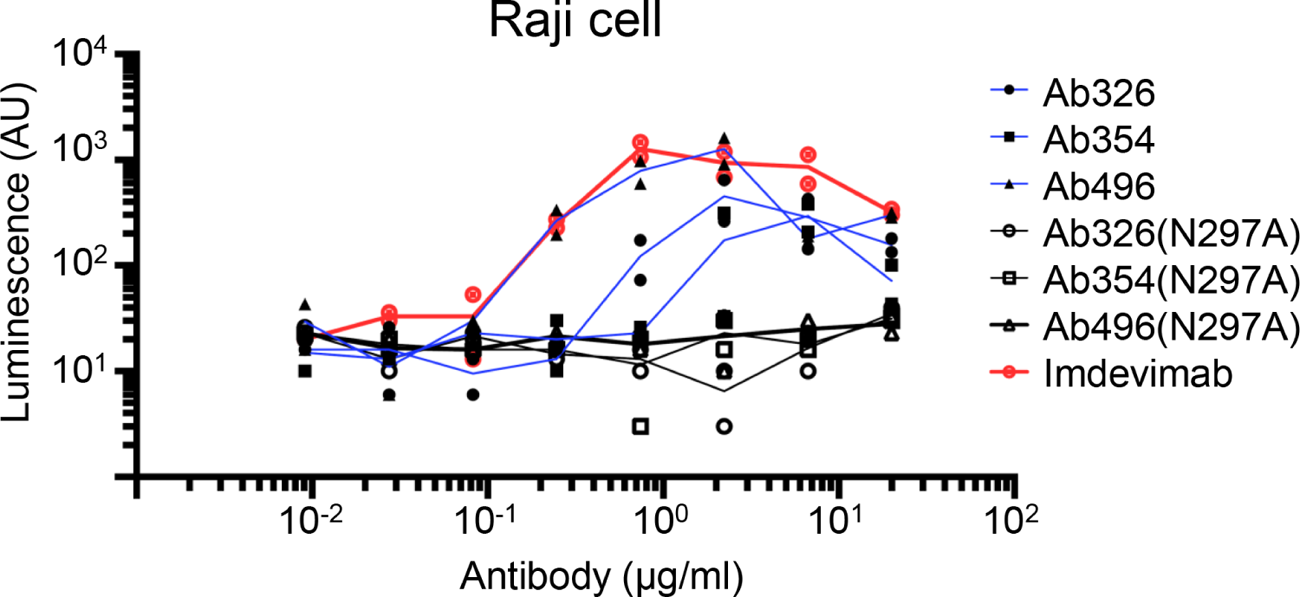
Fc-engineering for prevention of antibody-dependent enhancement Pseudovirus (Wuhan strain) was incubated with serially diluted antibody with or without N297A modification, and the mixture was applied to Raji cells. After incubation for 3 days at 37°C, the cells were lysed and subjected to luciferase assay. The lines represent mean value.

### *In vivo* treatment effect in two animal models

We next examined the *in vivo* effects of antibodies in two animal models, a hamster model (Imai et al., 2020) and a cynomolgus macaque model (Ishigaki et al., 2021). Hamsters were infected with the Wuhan strain on day 0, and were intraperitoneally treated with 50 mgkg BW of an N297A-modified antibody (Ab326, Ab354, or Ab496) or control human IgG1 on day 1. The dose of the antibodies was set at 50 mg/kg. Three days later, the amount of viral RNA in the lung tissue and the neutralizing antibody titer in the serum were measured (Figure 7A). Although antibodies could not be detected in the serum of some hamsters, probably due to technical issues during administration, viral RNA levels in lungs of the hamsters with sera that contained neutralizing antibody titers had reduced (Figure 7B).

**Figure 7.**
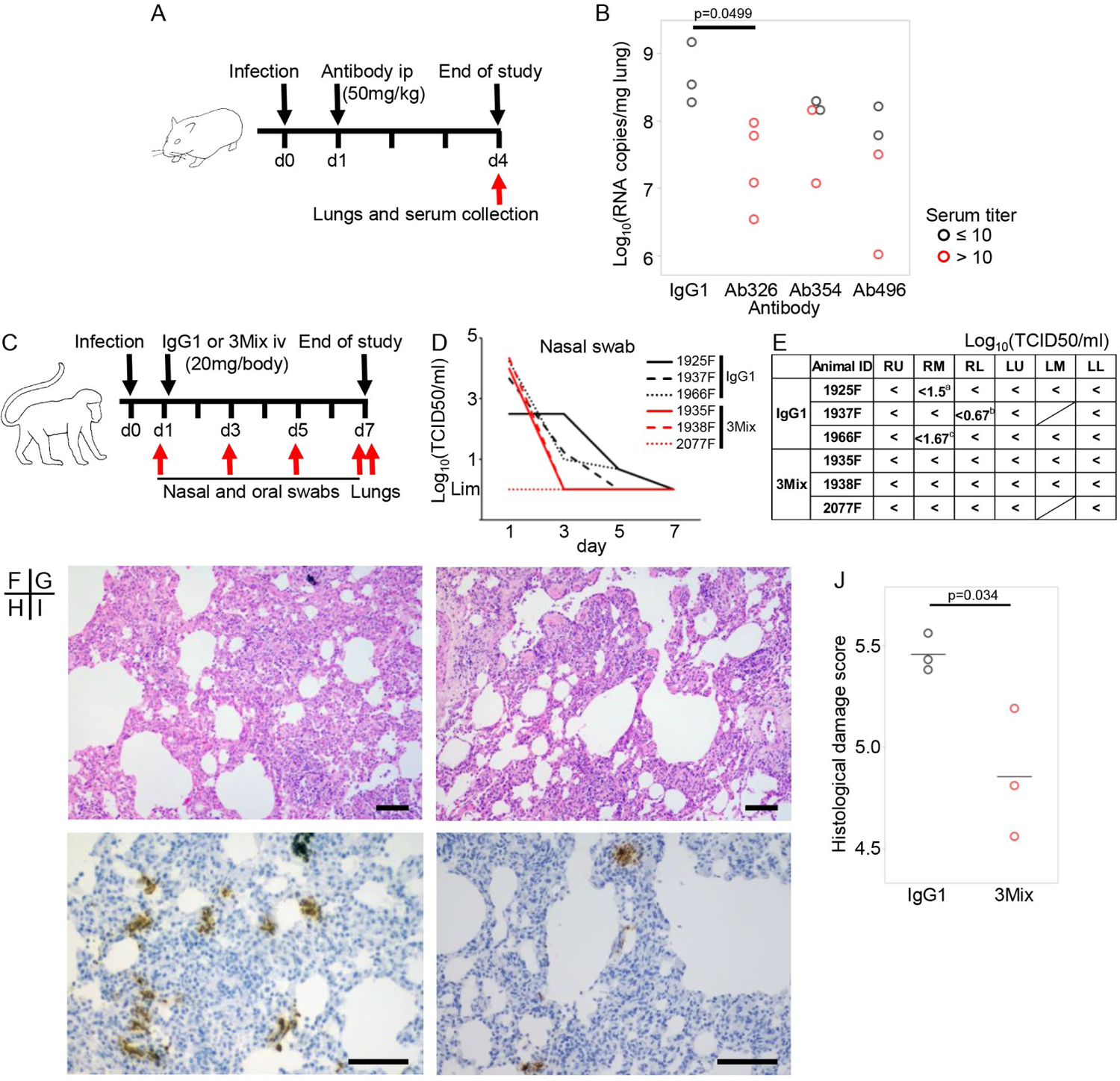
Therapeutic efficacy of neutralizing antibodies in two animal models (A) Overview of the experiment with the Syrian hamster model. Hamsters were inoculated intranasally with 10^3^ PFU of NCGM02. On day 1 post-infection, hamsters were injected intraperitoneally with 50 mg/kg BW of neutralizing antibodies or human IgG1 as a control. (B) On day 4 post-infection, serum and lung samples were collected. Serum neutralizing titer and viral RNA levels in the lung tissues were measured. Black circle indicates a low serum neutralization titer, and red circle indicates a high serum neutralization titer. \**p* < 0.05 by Dunnett’s test using all samples. (C) Overview of the experiment using the cynomolgus macaque model. Six cynomolgus macaques were inoculated with 2 × 10^7^ TCID_50_ SARS-CoV-2 JP/TY/WK-521/2020 into the conjunctiva, nasal cavity, oral cavity, and trachea on day 0. On day 1, 20 mg (5–7 mg/kg) of antibody cocktail 3Mix (1/3 each of an N297A-modified Ab326, Ab354, and Ab496) or human IgG1 as a control were injected intravenously. Nasal swabs were collected at day 1, 3, 5, and 7, and lung tissues were collected at day 7. (D) Viral titers of nasal swabs were measured. (E) Viral titers of each lung lobe were measured. “<” indicates under the detection limit. The shaded cells indicate that the lung lobe was not present. ^a^CPE-positive wells were 2/4 at no dilution, and 2/4 at 10^1^ dilution. ^b^CPE-positive wells were 1/4 at no dilution. ^c^CPE-positive wells were 1/4 at 10^1^ dilution. RU; right upper lobe, RM; right middle lobe, RL; right lower lobe, LU; left upper lobe, LM; left middle lobe, LL; left lower lobe. (F, G) Representative H&E stained sections and (H, I) immunohistochemical staining of SARS-CoV-2 N protein. (F, H) Lung sections of 1925F treated with the control antibody. (G, I) Lung sections of 1938F treated with the mixed SARS-CoV-2 specific neutralizing antibodies. The bars indicate 100 μm. (J) Histological damage score was evaluated according to the previously published criteria (Ogiwara et al., 2014). The bars indicate mean. \**p* < 0.05 by Student’s t test.

Finally, we examined the *in vivo* effects of antibodies in a cynomolgus macaque model. In this study, we used a cocktail of antibodies to cover broader mutations (VOCs had not yet appeared at the time of this study). We mixed equal amounts of Ab326, Ab354, and Ab496, and administrated a total of 20 mg/animal (5–7 mg/kg) the day after infection with a Wuhan strain. Nasal swabs were collected on day 1, 3, 5, and 7, and lung tissue samples were obtained on day 7 (Figure 7C). This model is known to spontaneously recover from COVID-19 in approximately 1 week (Ishigaki et al., 2021). Indeed, the live virus in the control IgG1 group became less than detectable range by day 5 to 7; however, those in all of the animals treated with mixture of the neutralizing antibodies became undetectable by day 3, earlier than the control group (Figure 7D). Finally, the lung tissues were evaluated by viral titer measurement and histological staining. Live virus was partially detected in the lungs of the control group even at day 7 whereas it was undetectable in the treatment group (Figure 7E). Interstitial pneumonia with lymphocytic infiltration and thickened alveolar walls were observed in the control macaques (Figure 7F) whereas such lesions were seen to a lesser extent in the macaques treated with the mixed neutralizing antibodies (Figure 7G). SARS-CoV-2 N protein–positive cell clusters were more frequently detected in the lungs of the control macaques than in the macaques treated with the mixed neutralizing antibodies (Figure 7H, I). Histological inflammation scores indicate that treatment with the mixed neutralizing antibodies reduced inflammation in the lungs, consistent with viral titers in the lung tissues (Figure 7J), without findings suggestive of ADE. These results demonstrated the *in vivo* antiviral effects of our antibodies.

## Discussion

We have produced many antibodies from the B cells of convalescent patients of Wuhan or D614G mutant strain, and obtained several neutralizing antibodies with high neutralization ability against variant strains. Initially, antibodies were produced from both antigen-nonspecific plasma cells and antigen-specific memory B cells, but the latter contained more superior antibodies, suggesting the importance of selecting B cells by antigen. The screening of neutralizing antibodies was performed by the cell-based Spike-ACE2 inhibition assay and cell fusion assay, which correlated with each other, and the results were confirmed by end-point authentic virus neutralization assay. Because antibodies were selected by competition with ACE2, antibody binding to Spike was classified as class 1/2, and epitopes on Spike were identified by cell-based mutated Spike-ACE2 inhibition assays and cryo-EM. Results from neutralization assays against Wuhan strain and VOCs using pseudoviruses and authentic viruses confirmed that selected antibodies were equivalent to or superior to imdevimab, which is used as a therapeutic agent. As for the *in vivo* function of these antibodies, efficacy for treatment use was demonstrated in both hamster and macaque models.

Although neutralizing antibodies are used as therapeutic agents and are highly effective, both bamlanivimab and casirivimab reduces susceptibility against E484 mutation (Starr et al., 2021), etesevimab reduces susceptibility against K417 mutation (Starr et al., 2021), and imdevimab reduces susceptibility against L452R mutation (Wang et al., 2021a). Most recently, the Omicron variant was demonstrated to reduce susceptibility of several therapeutic antibodies and antibody cocktails (Cameroni et al., 2021) (Planas et al., 2021). Oral therapeutic agents have also been rapidly developed, however, the fact that a single amino acid mutation in the viral neuraminidase resulted in resistance to neuraminidase inhibitors (Dapat et al., 2013; Lee and Hurt, 2018) insists on the importance to develop multiple therapeutics targeting various steps of SARS-CoV-2 infection.

One of the characteristics of our antibodies is N297 introduction on IgG1-Fc. This mutation almost eliminates binding to Fc receptors and, indeed, Fc-mediated uptake into Raji cells was eliminated. As for therapeutic antibodies, AZD7442 and etesevimab reduce binding to Fc receptors by YTE and TM modification (Loo et al., 2021) and LALA modification (Taylor et al., 2021) in the Fc domain, respectively, while bamlanivimab, imdevimab, and casirivimab are unmodified (Taylor et al., 2021). In contrast, sotrovimab increases binding to FcRn by LS modification (Gupta et al., 2021). In the absence of Fc receptor binding ability, there is a report that the therapeutic effect is decreased (Chan et al., 2021) and others that it is not significantly changed (Noy-Porat et al., 2021), and a consensus has not been established. In this context, our antibodies demonstrated the therapeutic effects in hamsters and macaques at doses of at least 5 to 7 mg/kg, without increased viral uptake by ADE (Figure 7).

In conclusion, we produced several Fc-modified neutralizing antibodies with high *in vitro* and *in vivo* efficacy. Because the virus is expected to continue to acquire mutations, it is desirable to prepare multiple therapeutic antibodies, and our antibody could be one of the candidates.

### Limitations of the study

This study has following limitations. The epitopes of our selected antibodies overlaps one another, and it is difficult to combine them as an antibody cocktail in which antibodies simultaneously bind their respective epitopes on a single molecule of Spike. The unevenly distributed epitopes of the selected antibodies are consistent with previous reports (Greaney et al., 2021; Piccoli et al., 2020) that antibodies in the sera of patients infected with Wuhan strains mainly target the area around E484. The number of animals that could be used was limited for both hamsters and macaques, although our antibodies were able to significantly reduce viral titers and tissue damage scores.

## Acknowledgments

We thank Yukari Kaneda, Mami Yamada, Mayumi Yonemochi, Mariko Ikeda, Mio Inoue, Naoko Kitagawa, and Hideaki Ishida for helping with the experiments, and Ikuo Kawamoto, Takahiro Nakagawa, Hideaki Tsuchiya, and Iori Itagaki for animal care. We also thank Dr. Takehiro Suzuki (Biomolecular Characterization Unit, RIKEN CSRS) for glycosylation analysis by mass spectrometry. The cryo-EM experiments were performed at the cryo-EM facility of the RIKEN Center for Biosystems Dynamics Research Yokohama. This study was supported by the Collaborative Research Resources, School of Medicine, Keio University, and was funded by a research grant for COVID-19 in Keio University School of Medicine (Donner Research Project Grant); by AMED under Grant Numbers JP20fk0108283, JP21ym0126022, and JP21fk0108468; by the RIKEN President’s discretionary funds; and by the Mitsubishi Foundation.

## Author contributions

M.T., H.S., Y.Kaneko, K.S. T.T. co-ordinated all of the research activities. M.T. wrote the initial draft of the manuscript. Y.T., M.S., K.K., and Y.I. contributed to the editing of the manuscript. H.S., M.Ishii, Y.Kondo, and K.F. contributed to sample collection. M.T. performed cell-based Spike-ACE2 inhibition assay, cell sorting and antibody production, biolayer interferometry. S.M. and Y.Takahashi performed authentic virus neutralization assay. H.F. performed fusion inhibition assay, pseudovirus/authentic virus neutralization assay, and viral uptake measurement by Fc receptors. C.M-O performed pseudovirus neutralization assay. M.S., K.K., T.U. performed cryo-EM analysis. T.Matsumoto, Y.Tomabechi., K.H. prepared recombinant antibodies and spike trimer protein. M.Imai, T.Maemura, Y.F., H.U., K.I-H., M.I., S.Y., Y.Kawaoka, and M.T. performed the studies in Syrian hamsters. M.T., H.I., M.N., C.T.N., Y.Kitagawa, and Y.I. performed infection study of cynomolgus macaques.

## STAR Methods

### RESOURCE AVAILABILITY

#### Lead contact

Requests for resources and reagents should be directed to the Lead Contact Author Masaru Takeshita.

#### Materials availability

All unique reagents generated in this study are available from the lead contact with a completed Material Transfer Agreements.

#### Data and code availability

The cryo-EM maps and atomic models have been deposited at the Electron Microscopy Data Bank and the PDB with the accession codes listed in Supplementary Table 3, and are publicly available as of the date of publication. Additional data needed to support the conclusion of this manuscript are included in the main text and supplementary materials.

### EXPERIMENTAL MODEL AND SUBJECT DETAILS

#### Ethics statements

We recruited 47 patients who had COVID-19, diagnosed by approved reverse transcriptase polymerase chain reaction (RT-PCR) tests for SARS-CoV-2 using swabs from the nose or saliva, and hospitalized at Keio University Hospital between April and December 2020. Peripheral blood samples from patients were collected at outpatient visits after discharge. The following parameters were collected from medical charts: signs and symptoms; lymphocyte counts; serum parameters of lactate dehydrogenase (LD), C-reactive protein (CRP); and medication history.

Disease severity is based on “COVID-19 Clinical Management,” edited by the World Health Organization (25 Jan 2021). This study was approved by the Ethics Committee of Keio University School of Medicine and conducted in compliance with the tenets of the Declaration of Helsinki. Informed consent was obtained from all participating individuals.

#### Animal studies

All experiments with hamsters were performed in accordance with the Science Council of Japan’s Guidelines for Proper Conduct of Animal Experiments. The protocols were approved by the Animal Experiment Committee of the Institute of Medical Science, the University of Tokyo. The study included one-month-old male hamsters.

All experiments with macaques were performed in accordance with the Guidelines for the Husbandry and Management of Laboratory Animals of the Research Center for Animal Life Science at Shiga University of Medical Science and Standards Relating to the Care and Fundamental Guidelines for Proper Conduct of Animal Experiments and Related Activities in Academic Research Institutions under the jurisdiction of the Ministry of Education, Culture, Sports, Science and Technology, Japan. The protocols were approved by the Shiga University of Medical Science Animal Experiment Committee. The study included eight- to nine-year-old female macaques.

#### Cells

293T and 293FT cells were maintained in DMEM containing 10% FCS and antibiotics at 37°C with 5% CO_2_. Expi293F cells were maintained in Expi293 Expression Medium at 37°C with 8% CO_2_. Vero E6/TMPRSS2 cells were maintained in low glucose DMEM containing 10% fetal bovine serum, antibiotics, and 1 mg/ml geneticin at 37°C with 5% CO_2_. Raji cells were maintained in RPMI1640 containing 10% FCS.

## METHOD DETAILS

### Production of vectors and proteins

The expression vectors and recombinant proteins were prepared as previously described (Takeshita et al., 2021a). Briefly, the extracellular domain of ACE2 (1-708 AA) was cloned into the pcDNA3.4 expression vector with a Flag tag or a streptavidin-binding peptide (SBP) tag at the C-terminus. Double-stranded DNAs coding full-length Spike of SARS-CoV-2 (UniProtKB-P0DTC2) and SARS-CoV-1 (UniProtKB-P59594) were codon-optimized and synthesized and inserted into the pcDNA3.4 vector. The extracellular domain (1-1213AA), S1 (1-685AA), and RBD (319-541AA) was cloned into the pcDNA3.4 vector with a SBP tag at the C-terminus. The expression vectors of mutated Spike were prepared by PCR using mutated primers. Recombinant RBD and ACE2 were produced using the Expi293 Expression System, and purified using Anti-DYKDDDDK tag Antibody Beads and Streptavidin Sepharose High Performance beads.

### Cell-based Spike-ACE2 inhibition assay

Cell-based Spike-ACE2 inhibition assay was performed as previously described (Takeshita et al., 2021a). Briefly, the expression vector of full-length Spike or mutated Spike and pMX-GFP were cotransfected into 293T cells using Polyethylenimine Max. After 2 days, cells were washed with PBS supplemented with 0.5% BSA and 2 mM EDTA (staining buffer), incubated with diluted serum samples or antibodies, washed again, and incubated with premixed ACE2-SBP and APC-conjugated streptavidin. After the final wash, the cells were analyzed by a FACS Verse. The median fluorescent intensity (MFI) among GFP^+^ cells was calculated using FlowJo and used as an indicator of Spike-ACE2 inhibition. The MFI of cells without serum or antibody were defined as 100% control and the ACE2-binding rate of serum or antibody was calculated as follows: ACE2-binding rate = 100 × (MFI of serum or antibody) / (MFI of control). Some serum results measured in a previous paper (Takeshita et al., 2021a) were used.

### Cell sorting

Peripheral blood mononuclear cells (PBMCs) were stored at −80°C and thawed using Anti-Aggregate Wash as previously described (Takeshita et al., 2019). CD19^+^ B cells were isolated by positive selection using CD19 Microbeads and stained with fluorochrome-conjugated antibodies and antigens for 20 min at 4°C. Plasma cells and antigen-specific memory B cells were defined as CD19^+^ surface IgD^-^ CD27^+^ CD20^-^ CD38^+^ and CD19^+^ surface IgD^-^ CD27^+^ CD20^+^ CD38^-^ Wuhan-RBD^+^ Wuhan-S1^+^, respectively, and sorted using a FACS Aria III flow cytometer.

### Production of antibodies

The production of single-cell cDNA was performed using Smart-seq2 (Picelli et al., 2014) with some modifications. Briefly, cells were sorted into 5 μl of Buffer RLT with 1% 2-mercaptoethanol in a 96-well PCR plate. RNAs were bound to 11 μl of RNAClean XP, washed using 80% ethanol as per the manufacturer’s instructions, and eluted by Elution buffer (2.3 μl of nuclease-free water, 1 μl of dNTP Mix, and 1 μl of modified oligo-dT primer. Subsequent reverse transcription PCR (23 cycles) and purification were performed as in Smart-seq2 (Picelli et al., 2014).

The production of expression vectors was performed as previously described with some modifications (Takeshita et al., 2020; Takeshita et al., 2021b). First, PCR was performed with a total volume of 20 μl containing 1 µl of cDNA library, 300 nM of primer set1, and 10 μl of KAPA HiFi HS ReadyMix. The cycling parameters were as follows: 95°C for 3 min; 30 cycles at 98°C for 20 s, 65°C for 15 s, and 72°C for 30 s; and 72°C for 1 min. For IgG, subsequently, second PCR was performed using primer set2 in a total volume of 20 μl containing 1 µl of 1^st^ PCR product, 300 nM of each primer, and 10 μl of KAPA HiFi HS ReadyMix with cycling parameters as follows: 95°C for 3 min; 30 cycles at 98°C for 20 s, 63°C for 15 s, and 72°C for 30 s; and 72 °C for 1 min. The IgG, Igκ, and Igλ products of second PCR were electrophoresed, purified, and inserted into the expression vector for IgH or Igκ or Igλ using NEBuilder HiFi DNA Assembly Master Mix.

When the second PCR failed to generate the appropriate band, third PCR was performed using primer set3 in a total volume of 10 μl containing 0.5 µl of 1^st^ PCR product, 300 nM of each primer, and 5 μl of KAPA HiFi HS ReadyMix with cycling parameters as second PCR. Third PCR products and second PCR products of IgA and IgM were sequenced. Fourth PCR was performed using primer set4 as in the second PCR, and the products were inserted into the expression vector. The primer list is shown in Supplementary Table 4. For *in vivo* studies, variable regions were inserted into the expression vector that had N297A mutation introduced in the Fc region.

Monoclonal antibodies were produced by transient cotransfection of IgH and IgL vectors using the Expi293 Expression System for 5 or 6 days, purified using Ab Capcher Mag2, and quantified using Human IgG ELISA Quantitation Set.

Anti-Spike protein antibodies for animal study were expressed using the Expi293 Expression System according to the manufacturer’s instructions. Culture media were harvested on day 7. The antibodies secreted were purified by Protein G Sepharose 4 Fast Flow resin, followed by gel-filtration chromatography on a Superdex200 10/300 GL column equilibrated in PBS.

### Biolayer interferometry

The affinity measurement of antibodies and epitope binning were performed by biolayer interferometry using Octet K2 and its software. For affinity measurement, anti-human IgG Fc Capture Biosensors were loaded by antibodies (20 μg/ml), exposed to a stepwise dilution of RBD solution (5.12–0.08 μg/ml) for 300 s, and then dissociations were observed in kinetics buffer for 300 s. The sensorgrams were adjusted by double references and fittings were performed with global analysis. For epitope binning, Streptavidin Biosensors were loaded by the extracellular domain of Spike (10 μg/ml) until the change in response reached 0.7 to 0.8 nm. After washing in kinetics buffer, the sensors were saturated with the first antibody (60 μg/ml) for 600 s. After confirming that the sensors were saturated by reacting 20 μg/ml of the first antibody, the sensors were reacted to 20 ug/ml of the second antibody to examine whether the two antibodies could bind simultaneously.

### Fusion inhibition assay

The assay was modified from the dual split protein (DSP) assay as previously described (Yamamoto et al., 2020). Two 293FT cell lines, one expressing SARS-CoV-2 Spike protein and DSP8-11, and the other expressing human ACE2, human TMPRSS2, and DSP1-7, were used in this assay. Both cells were treated with the final 6 μM EnduRen in the concentration of 2 × 10^5^/500 μl/well using 12-well plates. After making a single-cell suspension by pipetting, 50 μl (2 × 10^4^ cells) of the Spike-expressing cell suspension was incubated with 10 μl of 10 μg/ml of antibody for 30min at 37°C. Next, 50 μl (2 × 10^4^ cells) of the human ACE2/TMPRSS2 protein-expressing cell suspension was added to the antibody–cell mix and incubated for 2.5 h at 37°C. The Renilla luciferase activity was measured using the GloMax Discover System.

### End-point micro neutralization assay

End-point micro neutralization assay was performed as previously described (Matsuyama et al., 2020). Briefly, serially diluted sera were mixed with 100 TCID50 SARS-CoV-2 JPN/TY/WK-521 strain (Matsuyama et al., 2020) and incubated at 37°C for 1 h. The mixtures were placed on VeroE6/TMRRSS2 cells. After 5 days, plates were fixed with 20% formalin and stained with crystal violet solution. The highest sera dilution factor with 100% CPE inhibition was defined as the micro neutralization titer.

### Pseudovirus neutralization assay

Two plasmids, pNL4-3.luc.R-E- and pCMV3_SARS-cov2d19 series generated from the codon optimized pCMV3_Spike (1-1254AA) were transfected into cells and the pseudovirus was produced using the Expi293 expression system. Twenty thousand cells/100 μl/well human ACE2/TMPRSS2-expressing 293FT cell line (DSP1-7 (Yamamoto et al., 2020)) were seeded in a 96-well plate 1 day before infection. Serially diluted antibodies and 10,000 U/well viruses were mixed in 50 μl/well with DMEM containing 10% FCS, 1× MEM non-essential amino acids solution, and 1 mM sodium pyrubate, incubated for 37°C for 1 h. Fifty microliters per well of the cell culture supernatant were removed and the antibody–virus mixture was applied to the cell layer. After 48 h, the supernatant was removed completely and 50 μl/well of the Glo lysis buffer was added. After 5 min agitation, the plate was frozen at −80°C. After being thawed and agitated, 10 μl of the lysate was subjected to the Bright-GloTM Luciferase Assay system using a GloMax Discover microplate reader. The data from biological triplicates was analyzed using Prism 9 to determine the value of IC_50_ (ng/ml).

### Authentic virus neutralization assay

For Wuhan strain and Alpha, Beta, Gamma, and Kappa variant, eighteen thousand cells/well VeroE6/TMPRESS2 were seeded in a 96-well plate 1 day before infection. After the cell layers was washed with PBS, serially diluted antibodies and 100TCID50/well viruses were mixed in DMEM low glucose containing 2% FCS incubated for 37°C for 1 h, and applied to the cell layers. After 20 to 24 h, the cells were fixed, dried, and subjected to enzyme-linked immunosorbent assay (ELISA). The cells were washed with PBS containing 0.3% Tween-20 (v/v) as washing buffer three times and stained with rabbit anti-SARS-COV-2 nucleocapsid monoclonal antibodies in 5% Blocking One–containing washing buffer. After washing three times, the cells were incubated with HRP-conjugated donkey anti rabbit IgG(H+L) followed by washing and TMB colorimetric assay. Each sample was assayed in triplicate. All washing processes were performed using a 405 TS washer.

For Delta and Omicron (BA.1 and BA.2) variant, neutralization assays were performed as previously described {Moriyama, 2021 #118} with minor modifications. Briefly, a mixture of 100 TCID50 virus and serially diluted antibodies was incubated at 37C for 1 hour before being placed on VeroE6/TMPRSS2 cells seeded in 96-well flat-bottom plates. After culturing for 4-5 days at 37C supplied with 5% CO2, cells were fixed with 20% formalin and stained with crystal violet solution. Each sample was assayed in duplicate. Absorbance at 595 nm was measured using Epoch 2 area scanning mode and the neutralization percentage was calculated as follows; (sample signals – virus control signals)/(cell-only control signals – virus control signals) x 100. The data from biological triplicates were analyzed using Prism 9 to determine the value of IC_50_ (ng/ml).

### Viral uptake mediated by Fc receptors

Pseudovirus (Wuhan-hu-1 strain) was incubated with serially diluted antibody (3-fold; starting from 20 μg/ml) for 1 h at 37°C. The mixture of the antibody and the pseudovirus was applied to Raji cells. After incubating for 3 days at 37°C, the washed cells were lysed, and 10 μl of lysate was subjected to the luciferase assay.

### Production of proteins for cryo-EM

cDNA for the stabilized variant of SARS-CoV-2 Spike protein (residues 1-1208) with 6 beneficial proline substitutions (F817P/A892P/A899P/A942P/K986P/V987P) and “GSAS” substitution at the furin cleavage site (residues 682–685), named HexaPro (Hsieh et al., 2020), was subcloned into the pcDNA3.4 vector using PCR and In-Fusion Reaction. The above mutated Spike protein was designed to form a closed trimer by fusion with the C-terminal foldon trimerization motif followed by the TEV protease cleavage site, AviTag and 6xHis tag. cDNAs for heavy and light chains of the anti-Spike protein antibodies and their Fab heavy chain (VH-CH1) attached with the C-terminally TEV protease cleavage site and 6xHis tag were respectively subcloned downstream of the Ig kappa signal peptide into the pcDNA3.4 vector. Plasmid DNA was prepared using a GenElute HP Plasmid Maxiprep Kit.

Each protein was expressed using the Expi293 Expression System according to the manufacturer’s instructions. Culture media were harvested on day 4 for the Spike protein variant and day 7 for Fab. The Spike protein variant secreted in the culture medium was purified by a HisTrap HP column, followed by gel-filtration chromatography on a Superose 6 Increase10/300 GL column equilibrated in 20 mM Tris-HCl (pH8.0) and 150 mM NaCl. The secreted Fabs were purified with a HisTrap column, followed by gel-filtration chromatography on a Superdex200 10/300 GL column equilibrated in 20 mM Tris-HCl (pH8.0) and 150 mM NaCl.

Ab496 Fv-clasp was designed as described previously (Arimori et al., 2017) and cloned into a pCR2.1 TOPO TA cloning vector with an N-terminal His-tag and a TEV cleavage site fused to the VH and VL, respectively. Fv-clasp for Ab496 was synthesized at 25°C for 5 h in the presence of 5 mM GSSG and 0.4 mg/ml DsbC with the dialysis mode of the *Escherichia coli* cell-free protein synthesis method optimized to form disulfide bonds as described previously (Katsura et al., 2017; Matsuda et al., 2018). After the cell-free protein synthesis, the reaction solution was applied on a HisTrap HP column and the target protein was eluted with a linear gradient of imidazole. The His-tag was then cleaved by TEV protease and was removed by a HisTrap HP column. The protein was further purified by gel-filtration chromatography on a Superdex200 10/300 GL column in 20 mM Tris-HCl (pH8.0) and 150 mM NaCl.

### Cryo-EM data acquisition

The purified spike proteins (0.7mg/ml of the trimer) and each antibody were mixed with a molar ratio of 1:9, and incubated at 4°C for 30 min. The mixture (3 μl) was applied onto a freshly glow-discharged holey carbon grid, coated with a single or two layers of graphene oxide, then blotted for 3 s at 4°C in 100% humidity and plunge-frozen in liquid ethane using the Vitrobot Mark IV system.

Cryo-EM imaging of the complex was performed on a 300 kV Titan Krios G4 transmission electron microscope equipped with a BioQuantum K3 direct electron detector in the electron counting mode. The imaging was performed at a nominal magnification of ×105,000, corresponding to a calibrated pixel size of 0.829 Å/pixel. Each micrographic movie was recorded for a total of 15.5 electrons per pixel per second for 2.3 s, resulting in an accumulated exposure of 50.5 e-/A2. A total of ∼12000 movies were collected per data set. All of the data were automatically acquired using the EPU software, with a defocus range of −0.8 to −2.0 μm.

### Image processing

Image processing was mainly performed with the RELION-3.1.2 package (Zivanov et al., 2018). All of the dose-fractionated movies were subjected to beam-induced motion correction using MotionCor2 (Zheng et al., 2017), and the contrast transfer function (CTF) parameters were estimated using CTFFIND4 (Rohou and Grigorieff, 2015). Particles were picked using crYOLO (Wagner et al., 2019) from the micrographs and subjected to several rounds of 2D and 3D classifications to select good particles, and subsequently subjected to 3D refinement and Bayesian polishing. In the last cycle, particles were refined with a specific mask (imposing C3 symmetry on some complexes), yielding an overall map with a global resolution of 3.6 to 2.4 Å. To further improve the density of Fab moieties, the particles were subjected to focused 3D classification using a soft mask encompassing the RBD and variable domains of Fabs. The best-resolved and locally refined density maps were sharpened by the Phenix package (Liebschner et al., 2019). Reported resolutions are based on the Fourier shell correlation of 0.143 criterion.

### Model building and validation

The trimeric structures of the spike protein with each Fab were manually fitted and constructed based on a structure (PDB ID: 6VXX) as an initial model, using Coot (Emsley et al., 2010). The template model of the variable domain was automatically built by the SWISS-MODEL server (Waterhouse et al., 2018), based on selected sequences with higher homology, and was manually fit in Coot (Emsley et al., 2010). In the spike moiety, several prominent densities were identified near Asparagine residues, which corresponded to N-glycosylation. Glucosamines (NAG) were built depending on each density size. The model of the entire structure was iteratively refined using phenix.real_space_refine of the Phenix package (Liebschner et al., 2019), with secondary structure restrains. Model validation for stereochemistry was performed in MolProbity (Williams et al., 2018) (Supplemental table 3). Molecular graphics and density maps were prepared with PyMOL (https://pymol.org/2/) and UCSF ChimeraX (Goddard et al., 2018).

### Identification of *N*-glycan composition

The Ab354 antibody sample purified from HEK cells contains one *N*-glycosylation site at the Asn59 of the heavy chain, and the SDS-PAGE analysis revealed its heterogeneous modification. As previously described (Davydova et al., 2021), ∼1 nmol of the sample was digested by trypsin, and then treated with PNGase F to remove *N*-linked glycans as a control. Signals of glycopeptide from the positive fraction were detected using matrix-associated laser desorption/ionization time-of-flight (MALDI-TOF) mass spectrometry. Four major peaks were detected and subjected to estimate the glycan components by using Mascot. The four peaks were attributed to +dHex(3)Hex(4)HexNAc(5), +dHex(2)Hex(4)HexNAc(5), +dHex(1)Hex(4)HexNAc(5), and dHex(1)Hex(3)HexNAc(5). (To confirm the relative difference of each glycan unit, each fraction was further applied to LC-MS/MS.) Referring to the GlyConnect database (Mariethoz et al., 2018), possible glycan forms were listed (Supplementary Figure 2A). The traceable glycan model was reasonably placed near the Asn59 along the electron density, and fucose is identified, which is linked α-1,6 to the reducing terminal of β-*N*-acetylglucosamine, as shown in the Supplementary Figure 2BC.

### Experimental infection of Syrian hamsters

SARS-CoV-2/UT-NCGM02/Human/2020/Tokyo (Imai et al., 2020) was propagated in VeroE6 cells in Opti-MEM I containing 0.3% BSA and 1 µg of L-1-Tosylamide-2-phenylethyl chloromethyl ketone (TPCK)-trypsin/ml. One-month-old male Syrian hamsters (Japan SLC Inc., Shizuoka, Japan) were used in this study. Four hamsters per group were inoculated intranasally with 10^3^ PFU (in 100 μl) of NCGM02. On day 1 post-infection, hamsters were injected intraperitoneally with 50 mg/kg of Ab326, Ab354, Ab496, or human IgG1 Isotype control. The animals were euthanized on day 4 post-infection, and serum and lungs were harvested. Serum neutralizing titers were determined by using a plaque assay, and viral RNA copy numbers in the lung tissue were determined by RT-PCR.

For the neutralization assay, 35 µl of virus (140 tissue culture infectious dose 50) was incubated with 35 µl of twofold serial dilutions of serum for 1 h at room temperature, and 50 µl of the mixture was then added to confluent VeroE6/TMPRSS2 cells in 96-well plates, and incubated for 1 h at 37°C. After the addition of 50 µl of DMEM containing 5% FCS, the cells were incubated for 3 more days at 37°C. Viral cytopathic effects (CPEs) were observed under an inverted microscope and virus neutralization titers were determined as the reciprocal of the highest serum dilution that completely prevented CPEs.

For the RT-PCR, lung tissue was homogenized in 1 ml of DMEM containing 5% FCS and antibiotics. RNAs were extracted by using the QIAamp Viral RNA Mini Kit. RNA copies for the nucleoprotein (N protein)-encoding gene of SARS-CoV-2 were measured by using the TaqMan Fast Virus 1-step Master Mix. Amplification was carried out in 96-well plates on the StepOnePlus or CFX-96. The thermocycling conditions were as follows: 5 min at 50°C for reverse transcription, 20 s at 95°C for inactivation of reverse transcriptase and initial denaturation, and 45 cycles of 5 s at 95°C and 30 s at 60°C for amplification.

### Experimental infection of cynomolgus macaques

We performed *in vivo* infection assay using cynomolgus macaques as previously described (Ishigaki et al., 2021) with some modifications. Briefly, 2 × 10^7^ TCID_50_ WK-521 was inoculated into the conjunctiva, nasal cavity, oral cavity, and trachea of six cynomolgus macaques on day 0. On day 1, 20 mg (5–7 mg/kg) of antibody cocktail (1/3 each of Ab326, Ab354, and Ab496) or control human IgG1 was injected intravenously. Nasal swabs were collected at day 0 (before infection), day 1 (before injection), day 3, day 5, and day 7 (before sacrifice). Macaques were sacrificed at day 7, and lungs were collected.

To assess viral replication, serial dilutions of swab samples and tissue homogenate samples (10% w/v) were inoculated onto confluent VeroE6/TMPRSS2 cells for 1 h. The cells were washed with HBSS, and incubated with DMEM containing 0.1% BSA, penicillin, streptomycin, and gentamycin (50 μg/ml) for 3 days. CPEs were examined under a microscope. Virus titers were calculated by the Reed-Muench method.

To assess the viral amount at the nucleotide level, total RNA samples were extracted from swab samples using the QIAamp Viral RNA Mini Kit and RNeasy Plus Mini Kit according to the manufacturer’s instructions. RNA copies were measured as described above.

For histopathological examination, eight lung specimens were made from the bilateral lung of each macaque: one slice from each upper lobe and middle lobe, and two slices from each lower lobe. After fixing in 10% neutral buffered formalin for approximately 72 h, hematoxylin and eosin staining sections were made. Two pathologists blindly evaluated the histological score of the sections according to the previously published following criteria (Ogiwara et al., 2014): 0: normal lung, 1: mild destruction of bronchial epithelium, 2: mild peribronchiolar inflammation, 3: inflammation in the alveolar walls resulting in alveolar thickening, 4: mild alveolar injury accompanied by vascular injury, 5: moderate alveolar injury and vascular injury, 6, 7: severe alveolar injury with hyaline membrane-associated alveolar hemorrhage (under or over 50% of the section area). The average score of eight sections was calculated for each macaque, and the mean score of the two pathologists was defined as the histological score.

## QUANTIFICATION AND STATISTICAL ANALYSIS

Continuous data are presented as the median and interquartile range or as a number with the percentage value, as appropriate. Student’s *t*-test and Dunnett’s test were used to examine the continuous variables. Correlations between two continuous variables were analyzed using Spearman’s rank correlation coefficient. Unless stated otherwise, statistical analyses were performed using JMP. IC_50_ of each antibody were determined by non-linear regression using Prism 9, and shown in mean and SD. The sample sizes for the animal studies were determined from previous studies. The researchers were not blinded to the group allocations during the experiments. *p*-values < 0.05 were considered to be statistically significant.

### Supplementary file contents

Supplementary Table 1. The characteristics of patients screened for serum neutralizing ability

Supplementary Table 2. The characteristics of patients for antibody production

Supplementary Table 3. Cryo-EM data collection, refinement, and validation statistics for Spike RBD /Fab complexes.

Supplementary Table 4. Primer sequences

Supplementary Figure 1. Affinity and epitope binning Supplementary Figure 2.

Supplementary Figure 2. Identification of N-glycan composition

