## Supplementary file for "Fc-modified SARS-CoV-2 neutralizing antibodies with therapeutic effects in two animal models"

Supplementary Table 1. The characteristics of patients screened for serum neutralizing ability

| | Patients (n = 47) |
|---|---|
| Sex (male) | 25 (52.1) |
| Age (year) | 31.5 (27–55) |
| Smoking: never / ex / current | 35 / 0 / 13 |
| Days between first positive PCR test and first serum collection | 48.5 (36.5–58.5) |
| WHO classification[a] | Critical 2 (4.2)<br>Severe 1 (2.1)<br>Moderate 12 (25.0)<br>Mild 17 (35.4)<br>Asymptomatic 16 (33.3) |
| <Signs and symptoms> | |
| Fever (≥37.5°) | 26 (54.2) |
| Upper respiratory symptoms[b] | 24 (50.0) |
| Lower respiratory symptoms[c] | 18 (37.5) |
| Pneumonia | 15 (31.3) |
| <Laboratory data> | |
| Lowest lymphocyte count (/μl) | 1370 (967–1915) |
| Highest LD level (U/l) | 190 (158–235) |
| Highest CRP level (mg/dl) | 0.25 (0.03–1.3) |
| <Treatment> | |
| Systemic corticosteroids | 4 (8.3) |

Data are shown as numbers, numbers (%), or medians (interquartile ranges), as appropriate. [a]Disease severity is based on “Living guidance for clinical management of COVID-19,” edited by the WHO (23 Nov 2021). [b]Upper respiratory symptoms include rhinorrhea, sore throat, loss of smell, and loss of taste. [c]Lower respiratory symptoms include cough and sputum production.

WHO, World Health Organization; PCR, polymerase chain reaction; LD, lactate dehydrogenase; CRP, C-reactive protein

Supplementary Table 2. The characteristics of patients for antibody production

| Sample ID | Sex | Age | Days after first positive PCR | Severity | Fever | Pneumonia | Systemic steroid use |
|---|---|---|---|---|---|---|---|
| COV003 | male | 60 | 29 | severe | + | + | + |
| COV003-2 | male | 60 | 103 | | | | |
| COV011 | male | 65 | 39 | critical | + | + | + |
| COV011-2 | male | 65 | 60 | | | | |
| COV017 | female | 20 | 10 | moderate | + | + | - |
| COV019 | female | 30 | 55 | mild | + | - | - |
| COV025 | female | 36 | 44 | moderate | + | + | - |
| COV027 | female | 35 | 49 | mild | + | - | - |
| COV047 | male | 44 | 46 | mild | + | - | - |
| COV052 | male | 29 | 28 | moderate | + | + | - |
| COV080 | male | 33 | 62 | moderate | + | + | - |
| COV081 | male | 59 | 69 | moderate | + | + | - |
| COV082 | female | 65 | 53 | moderate | + | + | - |
| COV088 | male | 52 | 56 | critical | + | + | + |

PCR, polymerase chain reaction

Supplementary Table 3. Cryo-EM data collection, refinement, and validation statistics for Spike RBD /Fab complexes

| complexed with | Ab159 | Ab188 | Ab326 | Ab354 | Ab445 | Ab496(Fv-clasp) | |
|---|---|---|---|---|---|---|---|
| EMD ID | 33060 | 33061 | 33062 | 33059 | 33064 | 33063 | |
| PDB ID | 7X8Y | 7X8Z | 7X90 | 7X8W | 7X92 | 7X91 | |
| **Data collection and processing** | | | | | | | |
| Voltage (kV) | 300 | 300 | 300 | 300 | 300 | 300 | |
| Electron exposure ($e^-$/Å$^2$) | 50 | 50 | 50 | 50 | 50 | 50 | |
| Defocus range (μm) | | | -0.8 to -2.4 | | | | |
| Pixel size (Å) | 0.829 | 0.829 | 0.829 | 0.829 | 0.829 | 0.829 | |
| Symmetry imposed | *C*1 | *C*1 | *C*1 | *C*1 | *C*1 | *C*1 | |
| Particles in final reconstruction (no.) | 155607 | 69153 | 42808 | 74848 | 64299 | 79377 | |
| Map resolution (Å) postprocess | 4.1 | 4.1 | 4.2 | 3.1 | 4.1 | 4.3 | |
| FSC threshold | 0.143 | 0.143 | 0.143 | 0.143 | 0.143 | 0.143 | |
| Map sharpening B factor (Å$^2$) | -198 | -178 | -171 | -118 | -180 | -199 | |
| **Refinement** | | | | | | | |
| Initial model used | 7X8W | 7X8W | 7X8W | 5CCK | 7X8W | 7X8W | |
| RBD mode used | up | up | down | down | up | up | |
| Model resolution (Masked) (Å) | 4.3 | 4.3 | 4.3 | 3.3 | 4.3 | 4.4 | |
| FSC threshold | 0.5 | 0.5 | 0.5 | 0.5 | 0.5 | 0.5 | |
| **Model composition** | | | | | | | |
| Non-hydrogen atoms | 3435 | 3333 | 3301 | 3460 | 3358 | 3363 | |
| Protein residues | 440 | 426 | 420 | 435 | 426 | 427 | |
| Ligands | 1 | 1 | 2 | 8 | 2 | 2 | |
| B factors (Å$^2$) | | | | | | | |
| Protein | 111.02 | 85.93 | | 91 | 103 | 97 | 132 |
| Ligand | 88.78 | 96.08 | 108 | 128 | 114 | 130 | |
| R.m.s. deviations | | | | | | | |
| Bond lengths (Å) | 0.002 | 0.003 | 0.003 | 0.003 | 0.005 | 0.003 | |
| Bond angles (°) | 0.522 | 0.578 | 0.568 | 0.496 | 0.734 | | 0.615 |

| | | | | | | |
|---|---|---|---|---|---|---|
| **Validation** | | | | | | |
| MolProbity score | 1.89 | 2.16 | 2.11 | 1.61 | 2.26 | 2.13 |
| Clashscore | 6.67 | 10.46 | 9.9 | 4.44 | 12.5 | 11.08 |
| Poor rotamers (%) | 0 | 0 | 0 | 0 | 0.27 | 0 |
| Ramachandran plot | | | | | | |
| Favored (%) | 91.01 | 87.14 | 88.16 | 94.17 | 85.48 | 89.31 |
| Allowed (%) | 8.99 | 12.62 | 11.84 | 5.83 | 14.52 | 10.69 |
| Disallowed (%) | 0 | 0.24 | 0 | 0 | 0 | 0 |

Supplementary Table 4. Primer sequences

| | | 5' - 3' sequence |
|---|---|---|
| primer set1 | Forward | CAGTGGTATCAACGCAGAGTACG |
| | Reverse IgG | TGAGTTCCACGACACCGTCAC |
| | Reverse IgA | TTCGCTCCAGGTCACACTGAG |
| | Reverse IgM | ACAAAGTGATGGAGTCGGGAAG |
| | Reverse Ig κ | TCCGAGCTCGGTACCAAGCTAACACTCTCCCCTGTTGAAGCTC |
| | Reverse Ig λ | TCCGAGCTCGGTACCAAGCTATGAACATTCTGTAGGGGCCACT |
| | | TCCGAGCTCGGTACCAAGCTATGAACATTCCGTAGGGGCAAC |
| | | TCCGAGCTCGGTACCAAGCTATGAACATTCTGCAGGGGCC |
| primer set2 | Forward IgH | TGAGCTACGGACTCGAGCAGGTGCAGCTGGTGCAG |
| | | TGAGCTACGGACTCGAGGAGGTGCAGCTGGTGCAG |
| | | TGAGCTACGGACTCGAGGAGGTGCAGCTGGTGGAG |
| | | TGAGCTACGGACTCGAGGAGGTGCAGCTGTTGGAG |
| | | TGAGCTACGGACTCGAGCAGGTGCAGCTGCAGGAG |
| | | TGAGCTACGGACTCGAGCAGGTGCAGCTACAGCAGTG |
| | | TGAGCTACGGACTCGAGCAGGTTCAGCTGGTGCAG |
| | | TGAGCTACGGACTCGAGCAGGTCCAGCTGGTACAG |
| | | TGAGCTACGGACTCGAGCAGGTGCAGCTGGTGGAG |
| | | TGAGCTACGGACTCGAGGAAGTGCAGCTGGTGGAG |
| | | TGAGCTACGGACTCGAGCAGCTGCAGCTGCAGGAG |
| | | TGAGCTACGGACTCGAGCAGGTACAGCTGCAGCAG |
| | Forward Ig κ | TGAGCTACGGACTCGAGGACATCCAGATGACCCAGTC |
| | | TGAGCTACGGACTCGAGGACATCCAGTTGACCCAGTCT |
| | | TGAGCTACGGACTCGAGGCCATCCGGATGACCCA |
| | | TGAGCTACGGACTCGAGGATATTGTGATGACCCAGACTCC |
| | | TGAGCTACGGACTCGAGGATATTGTGATGACTCAGTCTCC |
| | | TGAGCTACGGACTCGAGGATGTTGTGATGACTCAGTCTCC |
| | | TGAGCTACGGACTCGAGGAAATTGTGTTGACACAGTCTCC |
| | | TGAGCTACGGACTCGAGGAAATAGTGATGACGCAGTCTCC |
| | | TGAGCTACGGACTCGAGGAAATTGTGTTGACGCAGTCT |
| | | TGAGCTACGGACTCGAGGACATCGTGATGACCCAGTC |
| | Reverse Ig λ | TGAGCTACGGACTCGAGCAGTCTGTGCTGACKCAGC |
| | | TGAGCTACGGACTCGAGCAGTCTGCCCTGACTCAGC |

| | | |
|---|---|---|
| | | TGAGCTACGGACTCGAGTCCTATGAGCTGACWCAGCC |
| | | TGAGCTACGGACTCGAGCAGCYTGTGCTGACTCAGTC |
| | | TGAGCTACGGACTCGAGAATTTTATGCTGACTCAGCCG |
| | | TGAGCTACGGACTCGAGCAGRCTGTGGTGACYCAG |
| | Reverse IgG | GATGGGCCCTTGGTGGA |
| | Reverse IgA | TCACACTGAGTGGCTCCTGG |
| | Reverse IgM | TCGTATCCGACGGGGAATTC |
| | Reverse Ig κ | GGAAGATGAAGACAGATGGTG |
| | Reverse Ig λ | AGCTCCTCAGAGGAGGGC |
| | | TCCTCAGAGGAGGGTGG |
| primer set3 | Forward IgH | ACAGGTGCCCACTCCCAGGTGCAG |
| | | AAGGTGTCCAGTGTGARGTGCAG |
| | | CCCAGATGGGTCCTGTCCCAGGTGCAG |
| | | CAAGGAGTCTGTTCCGAGGTGCAG |
| | Forward Ig κ | ATGAGGSTCCCYGCTCAGCTGCTGG |
| | | CTCTTCCTCCTGCTACTCTGGCTCCCAG |
| | | ATTTCTCTGTTGCTCTGGATCTCTG |
| | Reverse Ig λ | GGTCCTGGGCCCAGTCTGTGCTG |
| | | GGTCCTGGGCCCAGTCTGCCCTG |
| | | GCTCTGTGACCTCCTATGAGCTG |
| | | GGTCTCTCTCSCAGCYTGTGCTG |
| | | GTTCTTGGGCCAATTTTATGCTG |
| | | GGTCCAATTCYCAGGCTGTGGTG |
| | | GAGTGGATTCTCAGACTGTGGTG |
| | Reverse IgG | GATGGGCCCTTGGTGGA |
| | Reverse IgA | TCACACTGAGTGGCTCCTGG |
| | Reverse IgM | TCGTATCCGACGGGGAATTC |
| | Reverse Ig κ | GGAAGATGAAGACAGATGGTG |
| | Reverse Ig λ | AGCTCCTCAGAGGAGGGC |
| | | TCCTCAGAGGAGGGTGG |
| primer set4 | Forward | Optimized for each cell |
| | Reverse IgG | GATGGGCCCTTGGTGGA |
| | Reverse IgA | Optimized for each cell |
| | Reverse IgM | Optimized for each cell |

| | |
|---|---|
| Reverse Ig $\kappa$ | GGAAGATGAAGACAGATGGTG |
| Reverse Ig $\lambda$ | AGCTCCTCAGAGGAGGGC |
| | TCCTCAGAGGAGGGTGG |

Supplementary Figure

A

| | KD (M) | Kon (M/s) | Koff (/s) |
|---|---|---|---|
| **Ab287** | **<1.0E-12** | **3.01E+05** | **<1.0E-07** |
| **Ab326** | **5.14E-11** | **3.43E+05** | **1.76E-05** |
| **Ab354** | **7.26E-11** | **2.60E+05** | **1.89E-05** |
| **Ab376** | **1.10E-10** | **1.81E+05** | **1.98E-05** |
| **Ab496** | **<1.0E-12** | **6.95E+05** | **<1.0E-07** |
| **Ab336** | **2.35E-09** | **1.22E+05** | **2.86E-04** |
| **Ab159** | **1.24E-09** | **7.22E+05** | **8.95E-04** |
| **Ab175** | **<1.0E-12** | **9.63E+05** | **<1.0E-07** |
| **Ab188** | **<1.0E-12** | **2.60E+05** | **<1.0E-07** |
| **Ab308** | **6.62E-09** | **9.32E+04** | **6.16E-04** |
| **Ab445** | **5.93E-11** | **5.68E+05** | **3.37E-05** |
| **Ab302** | **<1.0E-12** | **1.53E+05** | **<1.0E-07** |
| **Ab315** | **<1.0E-12** | **1.72E+05** | **<1.0E-07** |
| **Ab369** | **1.97E-10** | **7.58E+04** | **1.50E-05** |
| **Ab176** | **<1.0E-12** | **8.31E+04** | **<1.0E-07** |
| **Ab374** | **<1.0E-12** | **1.17E+05** | **<1.0E-07** |
| **Ab114** | **<1.0E-12** | **2.12E+05** | **<1.0E-07** |

B

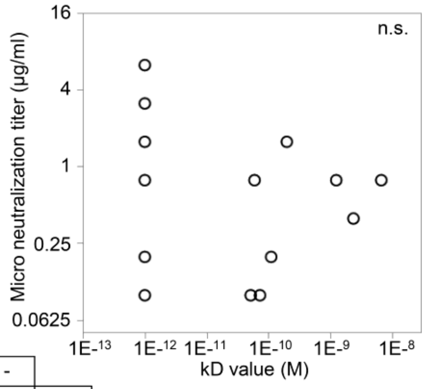

C

| | | | | | | | | | | | |
|---|---|---|---|---|---|---|---|---|---|---|---|
| **Ab287** | - | | | | | | | | | | |
| **Ab326** | - | - | | | | | | | | | |
| **Ab354** | - | - | - | | | | | | | | |
| **Ab376** | - | - | - | - | | | | | | | |
| **Ab496** | - | - | - | - | - | | | | | | |
| **Ab336** | - | - | - | - | - | - | | | | | |
| **Ab445** | - | - | - | - | - | - | - | | | | |
| **Ab159** | - | - | - | - | - | - | - | - | | | |
| **Ab175** | - | - | - | - | - | - | - | - | - | | |
| **Ab188** | - | - | - | - | - | - | - | - | - | - | |
| **Ab308** | - | - | - | - | - | - | - | - | - | - | - |
| | **Ab287** | **Ab326** | **Ab354** | **Ab376** | **Ab496** | **Ab336** | **Ab445** | **Ab159** | **Ab175** | **Ab188** | **Ab308** |

Supplementary Figure 1. Affinity and epitope binning

(A) The parameters of kinetics assays are shown. (B) The correlation of kD value and end-point micro neutralization titer is shown. Spearman’s rank correlation coefficient. n.s.: not significant. (C) The results of epitope binning are shown. The minus indicates overlapping epitopes.

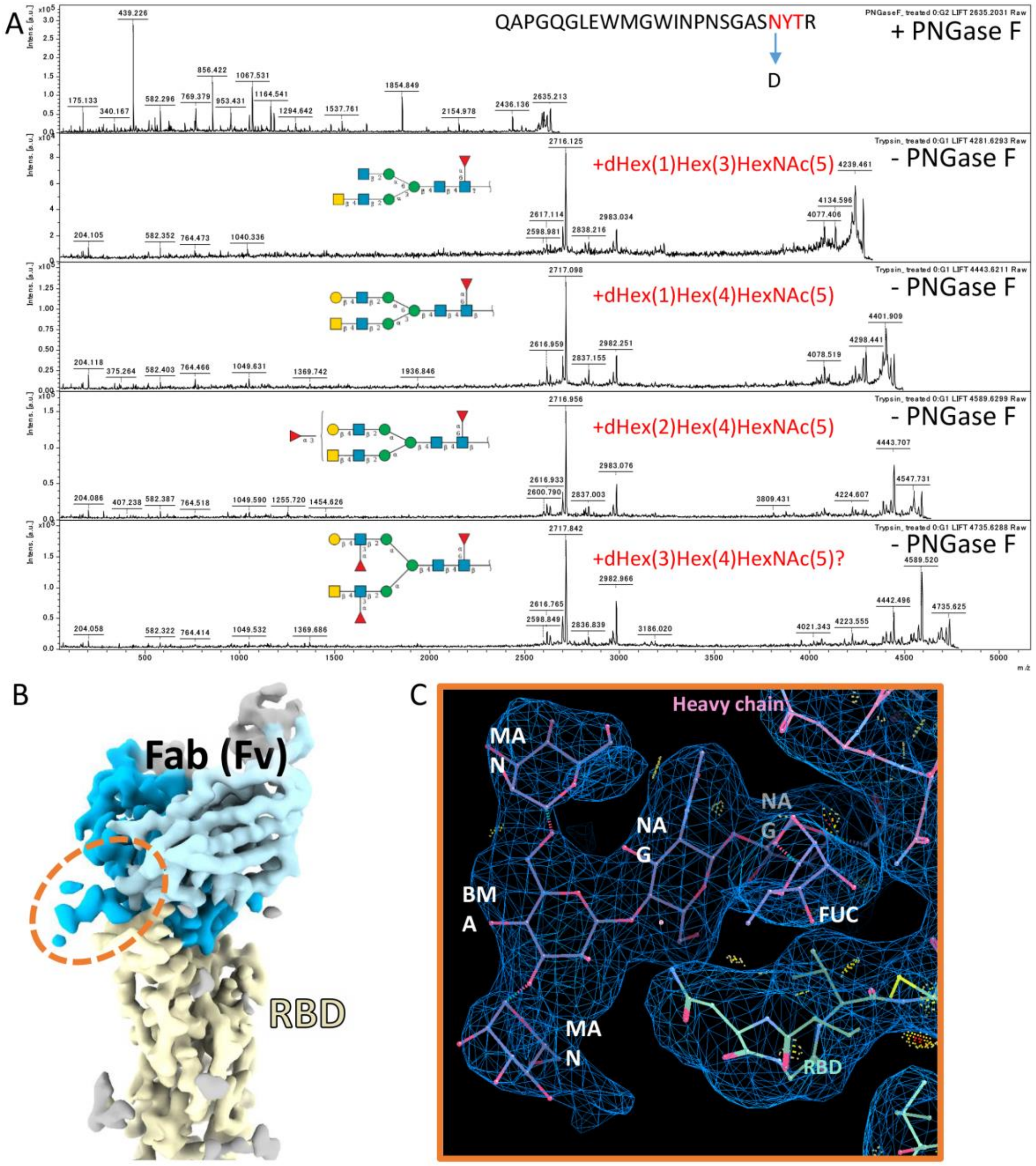

Supplementary Figure 2. Identification of N-glycan composition

(A) MS/MS spectrograms (300–5000 MW range) of the four candidates that contain a peptide of APGQGLEWMGWINPNSGASNYTR.

(B) Map after local refinement of RBD and the Fab region with specific mask. Dot circle shows the sugar density region of the heavy chain.

(C) The detail density of the sugar moiety with a traceable moiety.
